# The Influence of Irreversibly Electroporated Cells on Post Pulse Drug Delivery to Reversibly Electroporated Cells

**DOI:** 10.1101/2021.06.23.446669

**Authors:** Sid M. Becker

## Abstract

Electroporation can result in cell death in some proportion of a population of cells and, because the nature of the membrane disruption can vary significantly in irreversibly electroporated cells, there is uncertainty in the magnitude of and the transient behaviour of the associated permeability increases. This study numerically investigates the drug uptake by a population of cells that includes both reversibly and irreversibly electroporated cells. A theoretical continuum model is developed and simulations are conducted in conditions of low porosity (cells in tissues) and of high porosity (cells in suspension). This model estimates the permeability increases of electroporated cells using empirically based predictions of the dependence of long−lived electropore density on the local electric field magnitude. A parametric investigation investigates how the transmembrane transport characteristics of irreversibly electroporated cells (permeability and resealing rate) affect the drug uptake of the surviving cells. The results show that the magnitude and duration of the permeability increases of irreversibly electroporated cells is much more influential in low porosity tissues than in high porosity dilute suspensions. In applications of electroporation of cells in tissues, the uncertainty of irreversibly electroporated cells should be considered in regions of tissue experiencing field strengths for which the fraction of the total cells that are irreversibly electroporated exceeds about 0.1.

## 1 INTRODUCTION

Electroporation has been used in medical applications, such as electro-chemotherapy, DNA transfection for DNA vaccination and gene therapy, and tissue ablation [1–4]. The application of an intense electric field results in the introduction of nanometer-scale pores into the cell membrane [5–7]. Under certain conditions, these electropores can provide the sustained transient permeability increases that are required to facilitate the relatively slow diffusion process associated with post pulse transmembrane transport. Successful macroscale models of post pulse transport must reflect the physics involved these nm scale electropores; these include: the increases in cellular permeability associated with the introduction of the electropores, the pore resealing, any cell death associated with the electric field, and, if the electric field is to be modelled, changes in the bulk tissue electrical conductivity [8].

At moderate transmembrane voltages, reversible electroporation (RE) occurs in which transient electropores are introduced into the cell wall. Some of these pores are hydrophilic so that after the pulse, they experience a rapid transition back to the bi-layer membrane structure [5—7]. In some cases, the reversible electropores can be “long lived” lasting several minutes to hours after the pulse has been applied [9, 10]. Because this slow process of pore resealing can occur at a similar timescale as macroscopic diffusion, mass transport models that are dominated by the process of diffusion should include the effects of pore-resealing [8].

Irreversible electroporation (IRE) results when the effects of the electroporation are so extensive that the cell does not survive. Irreversible electroporation is reported to occur when the electric field experienced by a cell membrane exceeds a threshold value [11, 12]. In some instances, irreversible alterations of the cell membrane are formed during the first few seconds after electroporation and these result in sudden cell death; in other cases the cell membrane may not be irreversibly altered, though the cells continue to die for up to hours after the application of the final electroporation pulse. A theoretical model that attempts to predict the cellular uptake of drug by the surviving cells may also account for irreversible electroporation in regions of tissue exposed to extreme electric fields.

Experimental studies have shown that the magnitude of the increased permeability associated with electroporation increases with the magnitude of the applied electric field, the duration of the electroporation pulse, and the number of electroporation pulses delivered [12–14]. The extent of cell death associated with IRE has been shown to be similarly dependent on these pulse characteristics [11, 12]. Observations of populations of cells have shown the magnitude in the permeability increases and the extent of cell death follow sigmoidal dependences on the magnitude of the electric field [15], with the onset of increased permeability occurring at lower electric field magnitudes than those required for the onset of cell death. This is depicted in **Figure 1** which has been constructed to represent the data of [12].

**Figure 1.**
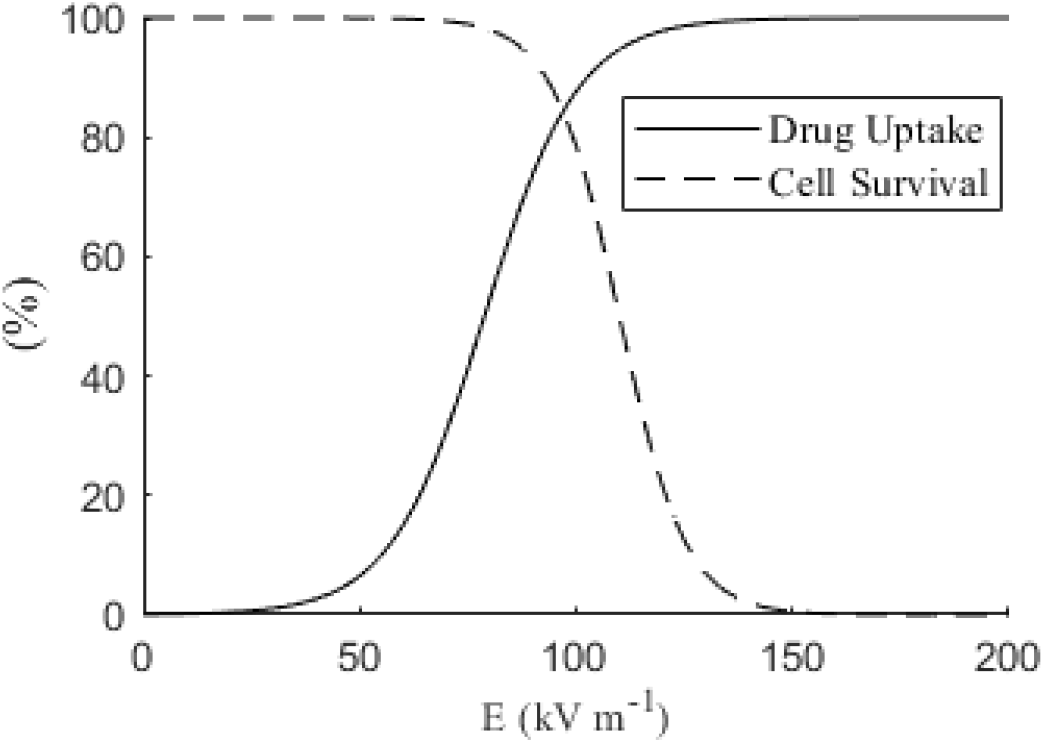
Cellular response to electric field magnitude: representative of experimental data presented in [12]. The solid line represents the percent of the surviving cells that experience an uptake of the drug. The dashed line represents the percent of the cells that survive the electroporation.

Different physics are attributed to different types of IRE cell death and these have been grouped into two general categories [11]. Sometimes the field results in sudden extensive and permanent damage to the cell membrane; here the associated electropores do not reseal. Sometimes the cell death is not attributed to such extensive membrane damage; here the electropores are both less extensive and they are temporary. In this case cell death is reported to be a result of “secondary” effects including: osmotic imbalances, loss of ATP, and the uptake of foreign molecules [11]. It is reasonable to anticipate that a cell membrane with large permanent defects will have a higher permeability to molecular transport than a cell membrane with smaller, temporary electropores.

The highlights of these are that for a particular cell type: (i) the degree of electroporation is field dependent, (ii) in the case of RE, the associated increases in permeability are not permanent, (iii) the cell death is field dependent, (iv) that significant cell permeability increases are observed electric field magnitudes that are lower than those required for irreversible electroporation, and (v) that cell death associated with irreversible electroporation may result from different physical mechanisms resulting in different transmembrane permeability characteristics. These points should be considered in theoretical continuum models of drug delivery to electroporated tissue and suspensions of cells.

One of the first developments in macroscale modelling of drug delivery in electroporated tissue is presented by Granot and Rubinsky in [16] in which the drug mass is conserved in a 2-compartment domain consisting of the extracellular space and the intracellular space. That study adapts the model of electroporation by Krassowska [5] to describe the influence of electric field magnitude on the state of the electropores (number and size of electropores per unit volume tissue). Granot and Rubinsky’s model includes cell resealing. Instead of calculating the permeability of the drug in the electropores, the study provides a parametric analysis of this parameter (over a one order of magnitude range). The Miklavčič group [17] extended this model by providing a first principles based description of the drug permeability in the electropores. The volume averaged drug permeability across the cell wall is estimated from the fraction of the total cell membrane area that is occupied by electropores. In a comparison with experiment, that study provides a parametric analysis of this fraction (over 5 orders of magnitude); so, this work does not explicitly estimate the permeability increases from the local electric field magnitude. In the study [8] a two-dimensional model is presented that includes field dependent changes in tissue’s electrical conductivity. Here the increase in the volume of cells electroporated is related to the local electric field magnitude, but the maximum permeability increases of the electroporated cells is not related to field strength. None of these studies distinguishes the drug delivered to the RE cells from the drug delivered to IRE cells (or that IRE and RE cells could have different permeabilities). The study [18] takes a step in this direction with a three equation model that uses a function curve fit from published data to relate the volume of tissue that is occupied by irreversibly electroporated cells to the magnitude of the local electric field (which is representative of the dashed line in **Figure 1**). Using a similar curve fit, that study also relates the volume of tissue occupied by cells that experience permeability increases to the magnitude of the local electric field (representative of the solid line in **Figure 1**). However that study does not relate the maximum permeability increases (for either RE or IRE cells) to the magnitude of the local electric field (which was implied in [16]); instead it assumes a constant value. That study also does not consider that IRE cells may experience resealing. None of the models reviewed fully relate the permeability increases to the local electric field magnitude and none are able to investigate the influence of the uptake characteristics of IRE cells on drug delivery to the RE cells.

In this paper a theoretical model is presented that is able to distinguish the drug uptake by RE cells from the drug uptake by IRE cells. The permeability increases of RE cells are explicitly related to the local electric field magnitude. The model is used in a zero-dimensional parametric study to investigate the whether the transport characteristics of IRE cells can significantly influence the drug delivered to RE cells, and if so, under what conditions.

## 2 THEORETICAL MODEL

The implications of Fig.1 are that electroporated populations of cells will be composed of cells that are reversibly electroporated as well as cells that are irreversibly electroporated. In the four equation model presented here, the drug mass is conserved in a domain that is composed of the extracellular space (EC) and the space occupied by three different types of cells. Irreversibly Electroporated (IRE) are the cells that do not survive electroporation. Reversibly Electroporated (RE) cells are surviving cells that are electroporated. For completeness we include Not Electroporated (NE) cells; these are surviving cells that do not exhibit permeability increases. This domain (**Figure 2**) represents the EC as the porous interconnected region that surrounds the cells. The cells are separated from one another by the EC so that there can be no direct cell to cell transfer of mass. In this study, the parameters of the domains are denoted by the subscripts: *EC*, *IRE*, *RE*, and *NE*.

**Figure 2.**
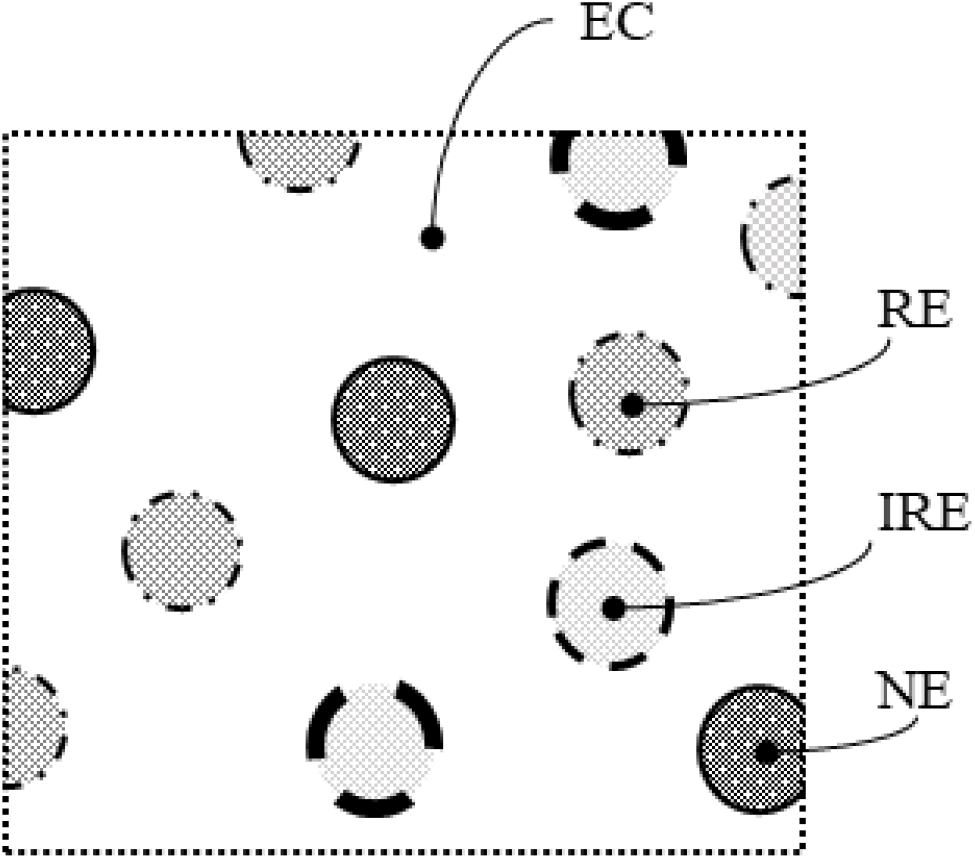
The 4 domains of electroporated tissue emphasizing the interconnectivity of the EC and the isolation of the individual cell types.

The porosity of the tissue is the ratio of the volume occupied by the extracellular space, V_*EC*_, to the total volume, V_*T*_:

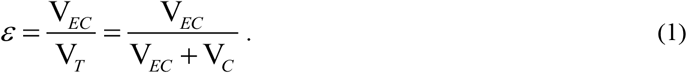

Here V_*C*_ is the volume occupied by all the cells; it comprises: (i) the volume occupied by irreversibly electroporated cells, V_*IRE*_, (ii) the volume occupied by the surviving cells that experience reversible electroporation, V_*RE*_, and (iii) the volume occupied by cells that do not experience permeability increases V_*NE*_:

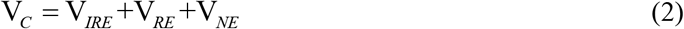

The fraction of the number of cells that are IRE (cells that do not survive) is represented by the parameter, *f_IRE_*. Under the approximation that the cellular volume is not influenced by the electroporation, the parameter is be related total cellular volume as:

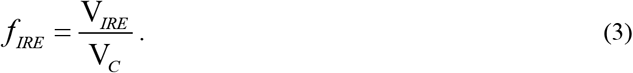

The fraction of the number of *surviving* cells that are permeabilized is represented by the symbol, *f _RE_*. Under the approximation that the cellular volume is not influenced by the electroporation, this parameter can be related to the volume of tissue occupied by the surviving cells by the relation:

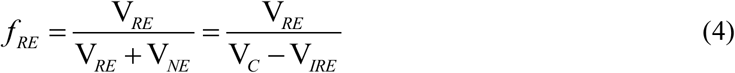

In this way the three volumes occupied by the three different cell types may be related to the total volume by:

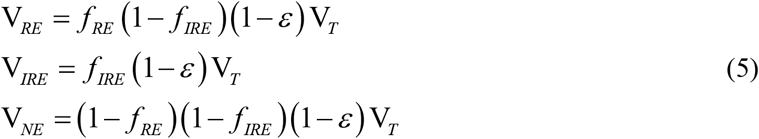

The intrinsic concentration is defined as the ratio of the mass stored in each sub-domain to the volume of that subdomain:

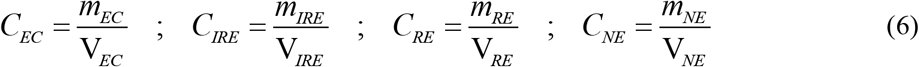

The intrinsic concentration differs from the total volume averaged concentration which is:

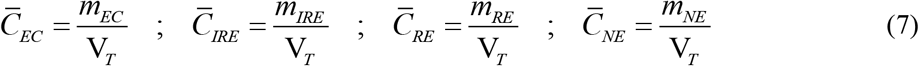

Substituting (5) and (6) into (7), the volume averaged concentrations may be related to the intrinsic concentrations by:

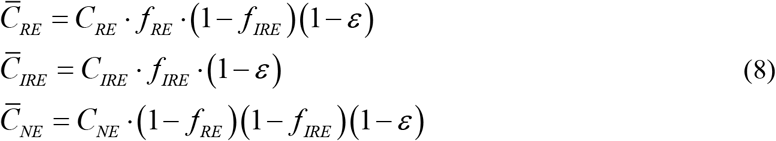

### 2.1 The General Governing Equations

A derivation of the set of equations governing mass transfer following the application of the electric field is provided in Appendix I. In the questions posed by this study, variations in spatial parameters are not considered, so that the diffusive term drops out and the PDE’s may be represented by a set of coupled ODE’s. Here the general expressions governing mass transfer in each domain are presented.

In the absence of spatial inhomogeneity, the intrinsic concentration of the drug in the extracellular space obeys:

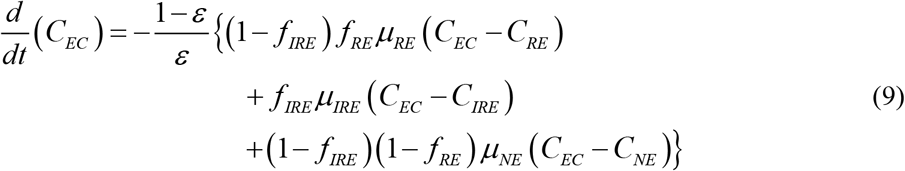

Here the subscripted symbols, *μ*, are the volumetric mass transfer coefficients (s^−1^) associated with the membranes of the different cells types. For IRE and RE cells, the magnitudes of these parameters should be representative of the effects of the electropores resulting from the applied electric field (which is discussed later).

The conservation of drug mass in the space occupied by surviving cells that do not experience permeability increases is governed by:

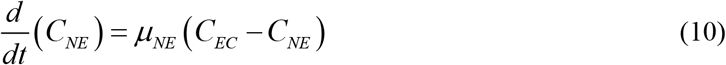

The conservation of drug mass in the space occupied by reversibly electroporated cells is governed by:

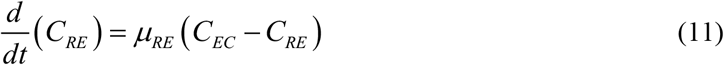

The conservation of drug mass in the space occupied by irreversibly electroporated cells is governed by:

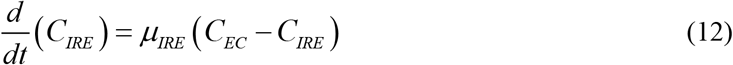

The parameters *μ*_RE_, *μ*_IRE_, *f _RE_*, and *f_IRE_* depend explicitly on the cell type and the electric field characteristics; this means that the magnitudes of these parameters should not be chosen independently or arbitrarily. The next sections address this dependence on electric field.

### 2.2 Field Dependence of Fraction of IRE Cells and of Fraction of Surviving RE Cells

Th electric field magnitude, the pulse duration, the number of pulses, and the pulse shape determine the extent of electroporation [5, 7, 9, 10, 12, 15, 19–27]. In some studies, the fraction of cells permeabilized under different conditions evaluated by counting the number of cells that exhibit evidence of increased permeabilization. In the study [15], the authors count the number of cells that die as a result of post pulse exposure to Bleomycin; the idea being that only the cells that were electroporated would take up the drug. The fraction of electroporated cells is expressed in that study as the ratio of the number of cells that die to the total number of cells that survive in the control group. In an earlier work [12], the authors count the number of cells that exhibit the uptake of fluorescent dyes at different electric field strengths and pulse durations. In that study for the dye, β-Gal, the permeabilization is expressed as the fraction of the cells that take up the dye to the total number of cells that survive (not the total number of cells). Both studies show a sigmoidal dependence of the fraction of the number of cells that are permeabilized (electroporated) on the applied electric field strength.

In [15] the authors explain this sigmoidal dependence; they note that at the individual cell level, (i) electropermeabilization is a result of the transmembrane potential difference, (ii) that this potential difference is proportional to the individual cell’s size and (iii) the size distribution of a population of cells is also sigmoidal. It is important to note that the fraction of cells permeabilized, *f _RE_*, is binary (the cells either are or they are not permeabilized). This fraction does address the degree of permeabilization: all cells electroporated will not have the same increases in permeability. In the studies of Refs. [12] and [15] and in the recent review by Jiang et al. in [11], the fraction of cells irreversibly electroporated, *f_IRE_*, (sometimes expressed instead as the survival fraction (*SF* = 1– *f_IRE_*) is shown to have a similar sigmoidal dependence on the electric field magnitude.

In the study [18] sigmoidal functions are curve fit to the data of Ref. [12] to predict the electric field dependence of the survival fraction and the fraction of cells permeabilized and those results are used here. For a particular pulse characteristic, the parameter *f RE* representing the fraction of the surviving cells that experience electropermeabilization may be related to the magnitude of the local electric field *E* (kVm^−1^):

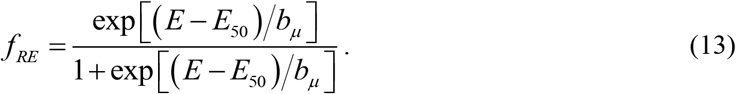

Here the constants *E*_50_ and *b_μ_* determine the shape of the sigmoid and their values may be determined using the experimental results. In comparison to the data of [12] using parameter values *E*_50_ = 78 kV·m^−1^ and *b_μ_* =10 kV·m^−1^ results in a mean absolute error of less than 3% (**Figure 3**a).

**Figure 3.**
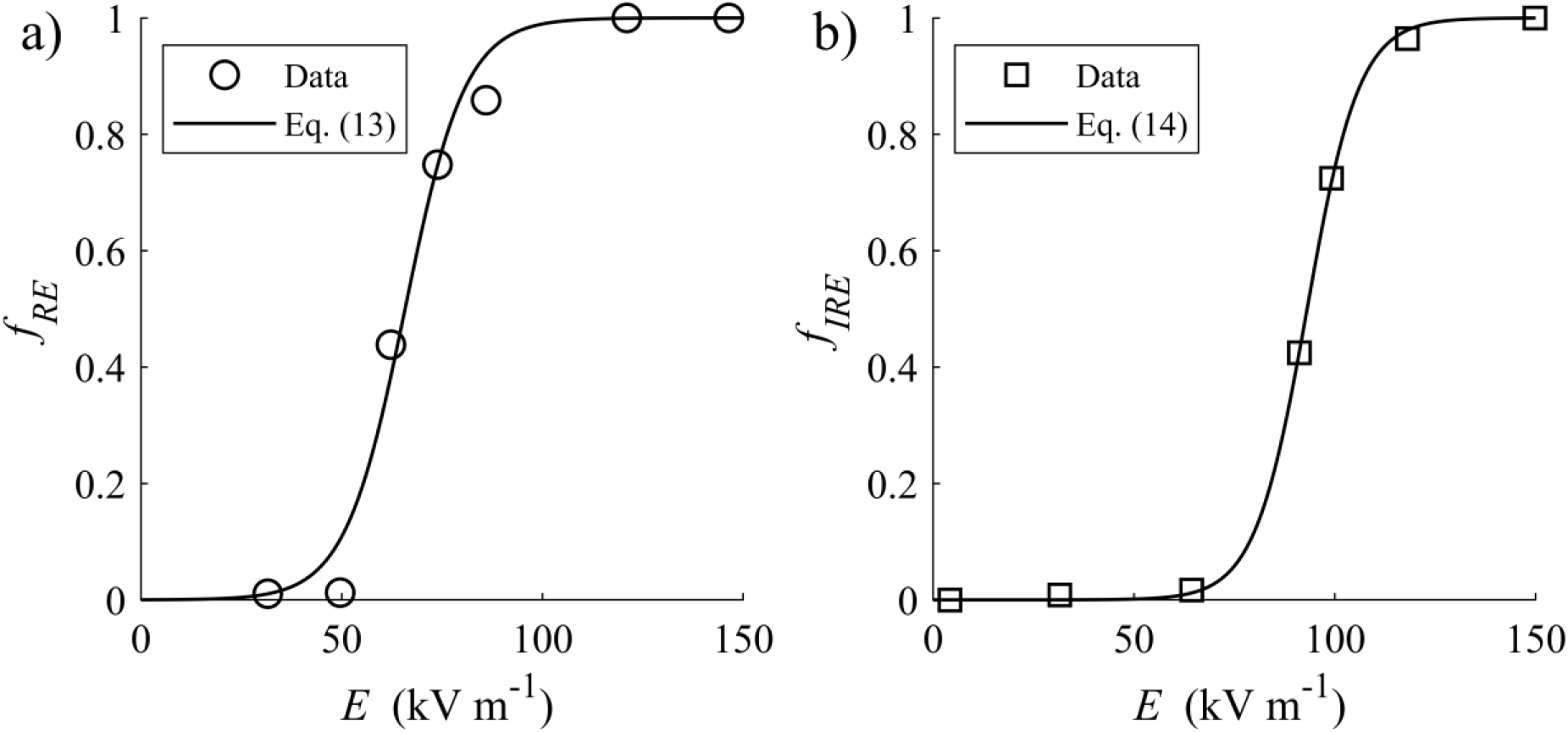
Comparison between published experimental data of [12] and the sigmoidal functions used in the current study taken from [18] for a) fraction of surviving cells permeabilized and b) fraction of total cells irreversibly electroporated.

The survival fraction of [18] is amended here to represent the fraction of cells that are irreversibly electroporated:

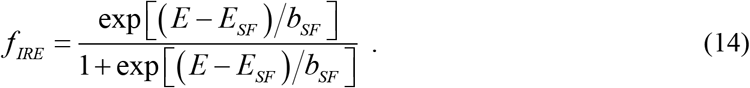

The prediction of Eq. (14) with the values, *E_SF_* =110 kV·m^−1^ and *b_sF_* = 7.9 kV·m^−1^, result in a mean absolute error of 1% compared to the data [12] (**Figure 3**b).

In any theoretical analysis electroporation of groups of cells, the values of parameters *f _RE_* and *f _IRE_* must not be chosen arbitrarily or independently of one another: both these parameters depend explicitly the cell type and the pulse conditions (the number of pulses delivered and the pulse field magnitude and duration).

### 2.3 Field Dependence of the Transient Mass Transport Coefficients

In this study the transport coefficients are used to represent the permeability of the different cell types: *μ_NE_* for the surviving cells that do not experience increased permeability, *μ_RE_* for the reversibly electroporated cells, and *μ_IRE_* for the irreversibly electroporated cells. These coefficients should satisfy the points:

(i) increases in permeability are associated with the introduction of the electropores into the cell wall, the number of electro-pores increases with electric field magnitude
(ii) that in RE cells (and perhaps in IRE cells) the changes in permeability are transient reflecting the resealing of the electropores

The NE cells are those that are unaffected by the application of the electric field so that in Eqs. (9) and (10), the mass transport coefficient *μ_NE_* is constant and is representative of the equilibrium permeability of the cells. This parameter depends on the drug characteristics and cell type and, in any practical application, its magnitude is expected to be much lower than those of electroporated cells; it is, after all, the goal of electroporation to overcome the limited transport associated with this parameter’s value.

The RE cells are known to experience increases in permeability with increasing electric field strength and as the pores reseal, the permeability values return to their pre-pulse values. Previous studies have represented the mass transfer coefficient of the RE cells with an exponentially decaying time dependence [8, 16, 18]:

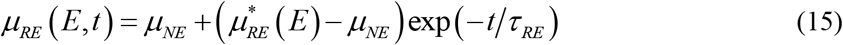

Here the mass transport coefficient 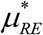 is representative of the average increase in permeability following electroporation of the cells that are reversibly electroporated. For an individual cell, the mass transfer coefficient is anticipated to increase with the electric field magnitude: this is because more pores are introduced into the cell walls at higher electric fields [5, 28, 29]. Briefly, in the theory of electroporation provided by [5], a mechanistic description underlying the evolution of the electropores is presented. During the pulse, the population of electropores have a range of sizes that can reach the order of 100 nm. However within 0.5 ms after the termination of the electric pulse, many of these disappear and any remaining electropores shrink to find a stable minimum energy state so that all remaining pores have a size of about 0.5 nm radius (which slowly reseal on a timescale of 10-100s) [30]. So if the post pulse electropores are uniform in pore size, the mass transfer coefficient associated with this state (immediately after the pulse) will be affected by the number of electropores present and this number is an effect of the pulse characteristics (number, duration, and magnitude). In [29] the authors show that the fraction of the total cell surface area that is occupied by these long lasting pores can be described by:

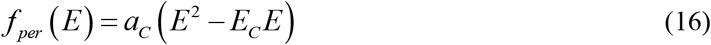

Here the constant *a_C_* depends on pulse amplitude, pulse duration, and pulse number and *E_C_* is the minimum electric field required for the formation of these electropores (often cited to be about 50 kVm^−1^). Fitting Eq. (16) to the electric field is discussed in the next section.

In the current study we follow the mechanistic description of the study [17] relating the increase in permeability of an individual cell to the fraction of cell membrane surface area occupied by these long lasting pores:

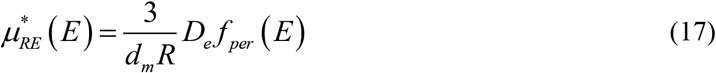

Here *d_m_* is the cell wall thickness and *R* is the cell radius. The term *D_e_* is the effective diffusivity of the drug in the electropores which is dependent on the drug’s hydrodynamic radius; this is described in the next section.

While the predictions of Eq. (17) have not been confirmed experimentally, it does provide a relative measure to estimate the order of magnitude of the permeability increases of reversibly electroporated cells. Regarding the mass transport coefficient associated with the IRE cells, there is much greater uncertainty. It is is possible that some IRE cells do not reseal at all (permanent pores) or that that they reseal over a similar timescale as those in reversibly electroporated cells. It is also possible that the IRE cells experience much larger maximum permeability increases (as in the case of the introduction of a large permanent pore). With such variability in mind, the IRE cell permeability changes will here be modeled to decrease exponentially in time; but the time constant and the maximum mass transfer coefficient may have different magnitudes than those associated with cells that are reversibly electroporated:

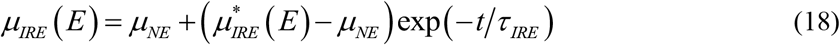

For the IRE cells, 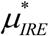 and *τ_IRE_* are more difficult to determine. Previous experimental studies have focused on the pulse conditions resulting in the cells’ binary response: survival (RE) or death (IRE). In such studies it would be of little interest to determine the rate of and the extent of a drug uptake to IRE cells following the application of the electric field. However, because it is believed that there are various mechanisms by which cell death occurs following electroporation [11], it can be anticipated that there may be great variability and uncertainty in the estimation of the mass transport coefficient increase, 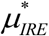, and the resealing time constant,*τ_IRE_*, associated with IRE cells. The influence of these parameters will be investigated in this study.

### 2.4 Parameter values

The rate at which the electropores reseal and the cell membrane is restored to its original state form is dictated by the time constant *τ_RE_*. This parameter is dependent on the pulse conditions, the cell and molecule type; in this study the value is assigned *τ_RE_* = 100 s^−1^ as in [18].

The field dependence of the increases in the mass transport coefficients of the RE cells are evaluated through Eqs. (15)–(17). The constant in Eq. (16) relating the fraction of the cell’s surface to the electric field was calculated from data taken from [29] for a train of 4 DC pulse of 100μs pulse duration applied at 1 s intervals (*f_per_* (*E*) = 3.4×10^−6^ at *E* = 86 kVm^−1^ with *E_C_* = 50 kVm^−1^) so that *a_C_* =1.186×10^−9^ m^4^kV^−2^.

Because of steric and hydrodynamic interactions between the solute (drug) and the pore wall, the effective diffusion coefficient appearing in Eq. (17) of the solute (drug) within the long lasting electropores will of course be much lower than the solute’s diffusion coefficient in water at low concentrations *D*_0_. Many models estimating this resistance to diffusive transport in cylindrical pores use a simple hindrance analogy [31]:

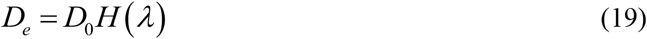

Here the hindrance factor, *H*, is characterized by the ratio of the solute hydrodynamic radius to the electropores radius, *λ* = *r_S_/r_P_*. An accurate expression for the range 0≤*λ*≤ 0.95 is [31]:

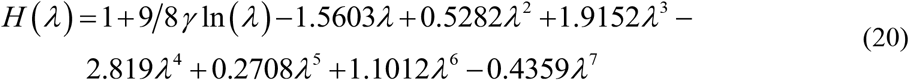

The permeability increases of the RE cells are evaluated with parameter values close to those of [17]; these are representative of sucrose: *D*_0_ = 4.5×10^−10^ m^2^s^−1^, *d_m_* = 5×10^−9^ m, *R* = 2.5×10^−5^ m and *λ* = 0.8. For these conditions, the dependence of the permeability increases of the RE cells, 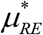, on the electric field magnitude is plotted in **Figure 4** which reflects the fractional pore area with increasing electric field reported in [29]. The parameter values of this study are listed with citations in Table 1.

**Table 1.** Parameter values used in the study.

| Parameter | Symbol and value | Used in Eq. |
| --- | --- | --- |
| Constants of fraction of surviving cells electroporated | $E_{50} = 78 \text{ kV} \cdot \text{m}^{-1}$, $b_{\mu} = 10 \text{ kV} \cdot \text{m}^{-1}$ [18] | (13) |
| Constants of fraction of total cells not surviving | $E_{SF} = 110 \text{ kV} \cdot \text{m}^{-1}$<br>$b_{SF} = 7.9 \text{ kV} \cdot \text{m}^{-1}$ [18] | (14) |
| Field magnitude at onset of electroporation | $E_C = 50 \text{ kVm}^{-1}$ [29] | (16) |
| Constant in fraction cell membrane area occupied by pores | $a_C = 1.186 \times 10^{-9} \text{ m}^4 \text{ kV}^{-2}$ (calculated) [29] | (16) |
| Thickness of cell membrane | $d_m = 5 \times 10^{-9} \text{ m}$ [17] | (17) |
| Average cell radius | $R = 2.5 \times 10^{-5} \text{ m}$ [17] | (17) |
| Diffusivity of sucrose in water | $D_0 = 4.5 \times 10^{-10} \text{ m}^2 \text{ s}^{-1}$ [17] | (19) |
| Ratio: drug hydrodynamic radius to electropore radius | $\lambda = 0.8$ similar to [17] | (20) |
| mass transfer coefficient of NE cells | $\mu_{NE} = 10^{-7} \text{ s}^{-1}$ (estimated) | (21) |
| RE cell electropore resealing time constant | $\tau_{RE} = 100 \text{ s}^{-1}$ [18] | (21) |

**Figure 4.**
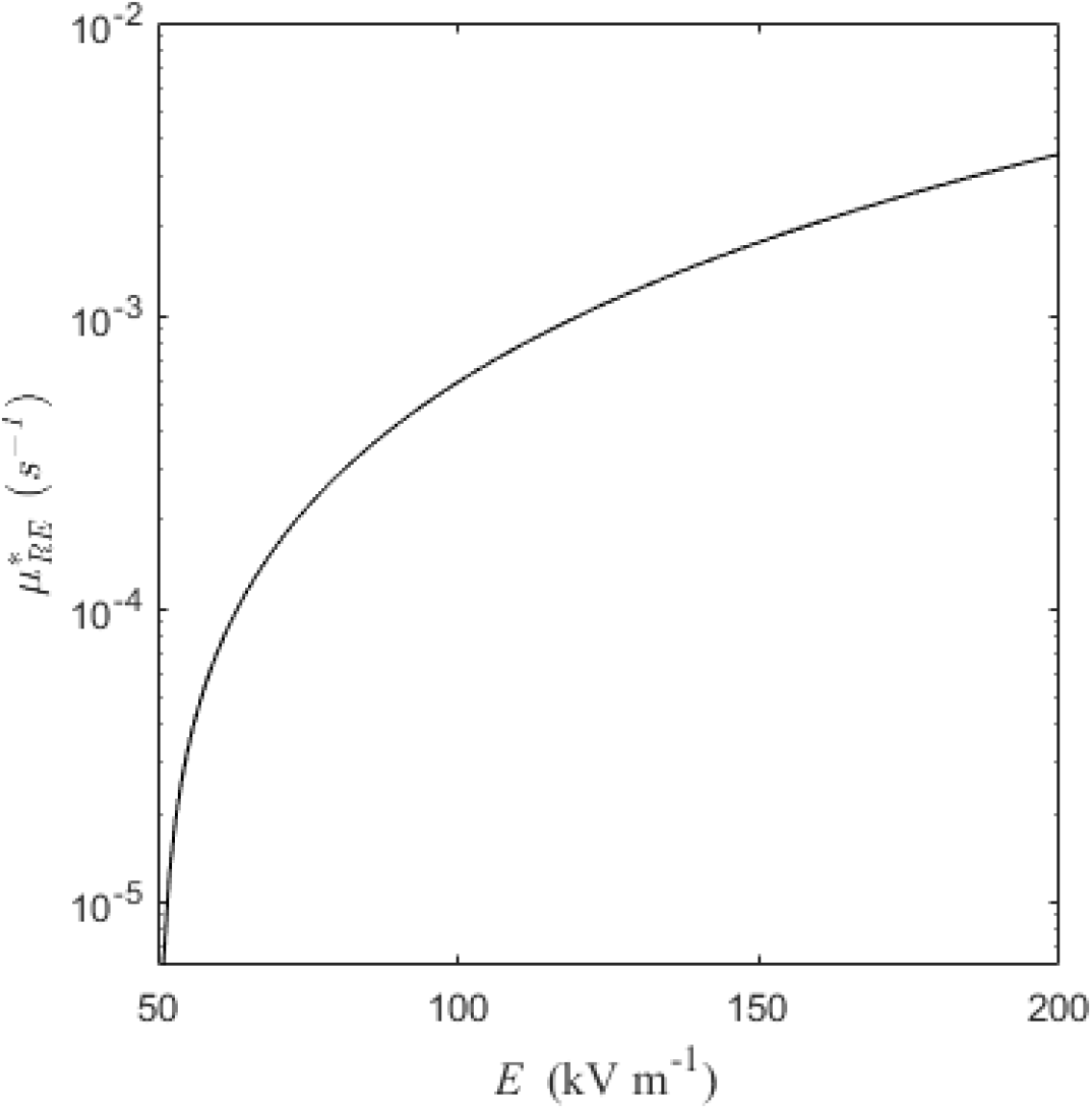
Field dependence of the permeability increases resulting from Eqs. (17), (19), and (20) with parameter values of **Table 1**.

The mass transport coefficient of non-electroporated cells is anticipated to be very small so that in this study it is assigned a value of *μ_NE_* =10^−7^ s^−1^, which is 2 orders of magnitude below the RE permeability increases at low field strength (**Figure 4**). The transport related parameters of the IRE cells have been chosen to represent two different causes of cell death. One reflects the immediate cell death resulting from the complete destruction of the cell membrane; here very large and permanent pores are introduced to the cell membrane. In the other case, cell death is a result of osmotic imbalances associated with smaller pores that reseal. To reflect these conditions, parametric investigations are conducted on the parameters *τ _IRE_* and *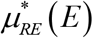*. Two values of the time constant *τ _IRE_* were considered. In one, the IRE cells reseal at the same rate as the RE cells (*τ _IRE_* = *τ _IRE_*). To represent the case in which the cell wall is permanently damaged (*τ _IRE_* → ∞) a very large value is assigned: *τ _IRE_* =10^10^s. Two magnitudes of IRE permeability increases are considered. It seems reasonable to consider that the IRE cells would not have permeability values below those of the surviving cells, thus the minimum value of the increase in transport coefficient associated with the IRE cells was chosen to be equal to that of the RE cells 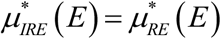. As an upper limit to the IRE cell permeability, to represent the case when the cell wall is completely destroyed, a value 100 times larger than that of the reversibly electroporated cells was assigned 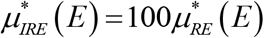.

In the parametric investigations that follow, two different values of the porosity (reflecting cell density) are chosen. A high porosity *ɛ* = 0.8 is used to represent the electroporation of a dilute suspension of cells. To represent the high density of cells in tissue, a low value of porosity of *ɛ* = 0.2 is chosen (similar to the reported porosity of liver [32]).

### 2.5 Numerical Modelling

For clarity, the complete set of ODE’s that are numerically evaluated are presented here in their full form:

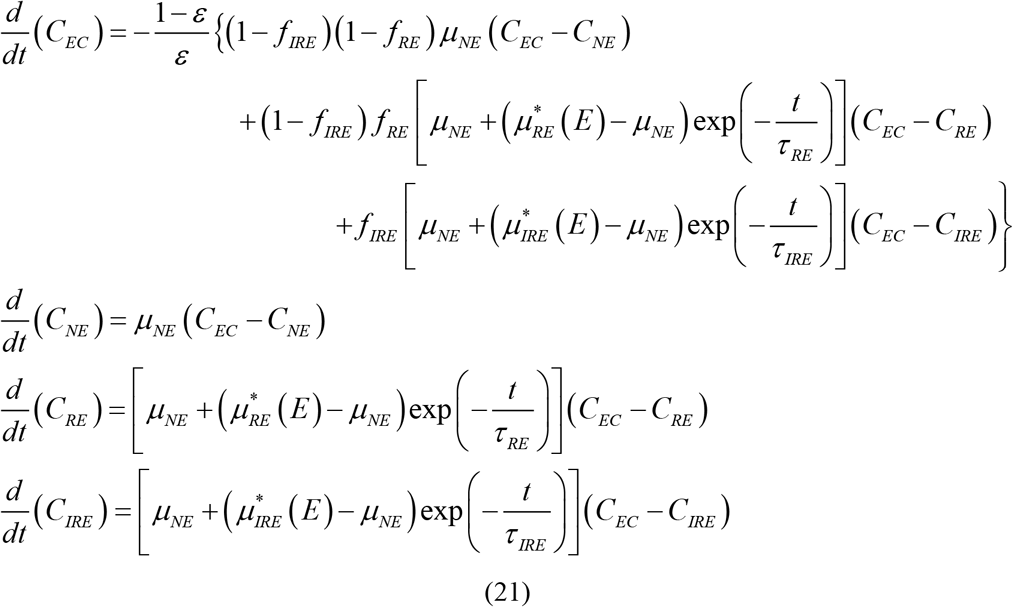

In system of Eqs. (21) the volume fractions *f _RE_* and *f _IRE_* are related to the electric field by Eqs. (13)–(14) and the mass transport coefficient 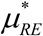 is evaluated with Eqs. (17), (19)–(20) using the parameter values presented in Table 1. These equations are approximated numerically using the MatLab explicit Runge-Kutta time integration solver, ode45. Relative tolerances of 10^−9^ were used. The initial intrinsic concentrations in the extracellular space and in the spaces occupied by the three cell types were set to:

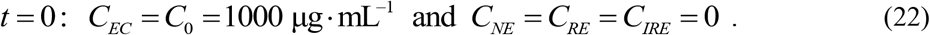

Time integration at regular 0.1s time steps was conducted for the first 400s. As a means of verifying the results of this numerical integration, a comparison with the analytic solution to a similar (but much simplified) problem is developed and shown in Appendix II.

## 3 Results and Discussion

The motivation behind the development of the system of ODEs of Eq. (21) is to evaluate the efficacy of the delivery of drug to the surviving RE cells (*C_RE_*). The uncertainty underlying the irreversibly electroporated cells’ parameter values *μ_IRE_* and *τ _IRE_* hinder the confident prediction of the drug uptake by reversibly electroporated cells. This study presents a parametric investigation of the influence of these parameters on *C_RE_* for different porosity cases: a low porosity case reflecting the conditions representative to cells in tissue and a high porosity case representative to cells in suspension.

The drug uptake by the cells in the 400 s after the final pulse at different field magnitudes is considered first (**Figure 5**). At intermediate field strengths, the influence of IRE cell properties on RE cell uptake is negligible: less than 1% difference for the low porosity case at field strengths below 67 kVm^−1^ and for the high porosity case at field strengths below 88 kVm^−1^. At an electric field strength of 75 kVm^−1^ only about 1% of the cells have been irreversibly electroporated (the top horizontal axes of **Figure 5** map the fraction of cells irreversibly electroporated to the field strength). At higher field strengths, as the fraction of cells irreversibly electroporated increases, so does their influence on the intrinsic RE cell concentrations. These influences are more noticeable in the low porosity case **Figure 5**a than in the high porosity case **Figure 5**b).

**Figure 5.**
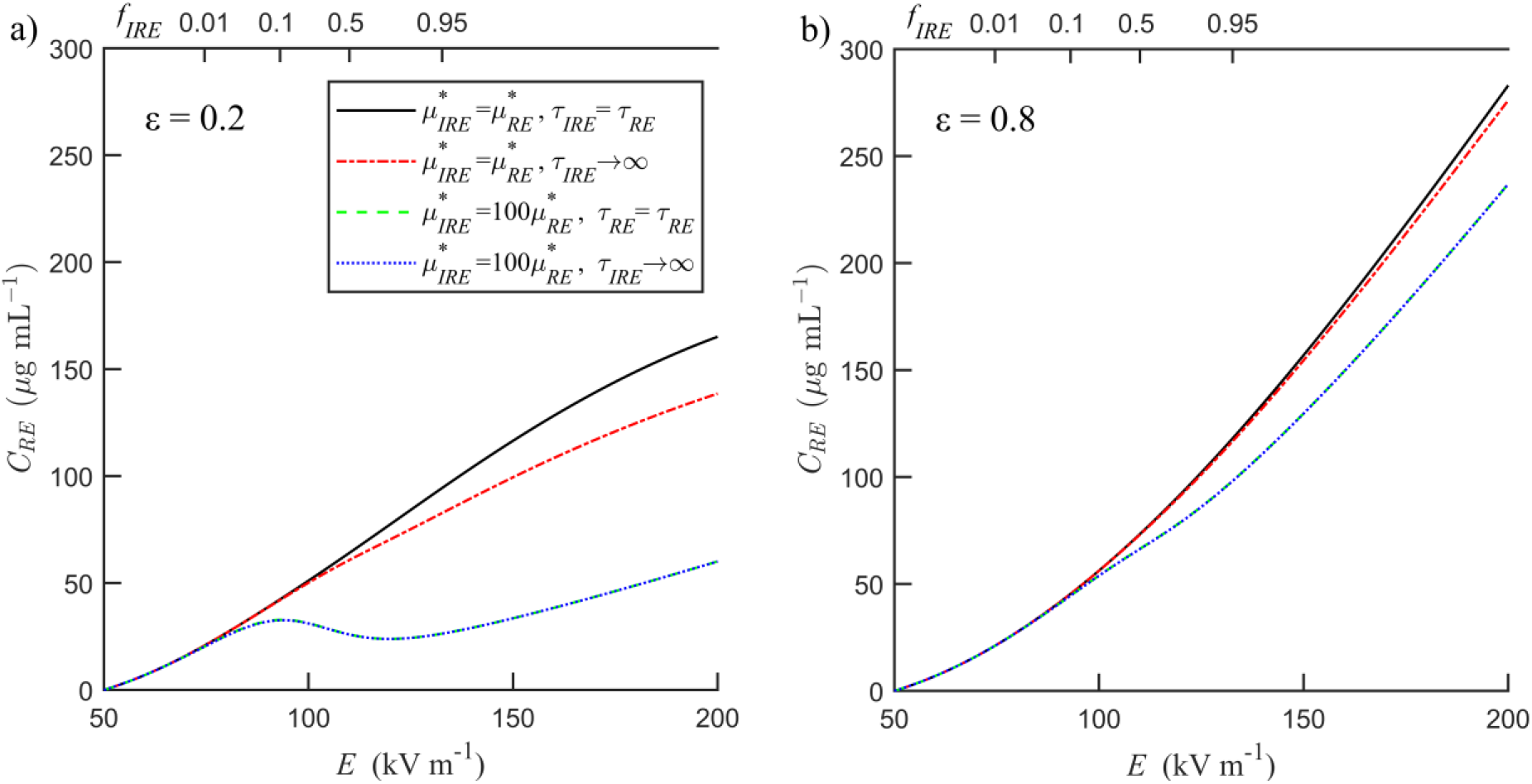
Field dependence of intrinsic concentrations in the space occupied by reversibly electroporated cells at 400 s post pulse for a) low porosity case, b) high porosity case. The fraction of total cells that are irreversibly electroporated is expressed on the top axis.

Setting the IRE permeability increase equal to that of RE cells 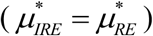 reflects cell death that result from “small pore area” osmotic imbalances. Here the influence of IRE cell resealing (represented by the exponential decay term,*τ _IRE_*) is seen in the solid black line and the dash-dot red lines of **Figure 5**a,b. For the low porosity condition (reflecting cells in tissue, **Figure 5**a) at field strengths below 95 kVm^−1^ the percent difference in drug uptake to RE cells (with and without IRE resealing) is less than 1% ; this difference increases with field strength (until it reaches nearly a 20%). In the high porosity case condition (reflecting cells in dilute suspensions, **Figure 5**b), the influence of cell resealing on drug uptake to RE cells is negligible: there is a less than 1% difference in concentrations at field strengths below 127 kVm^−1^ and even at the upper field strength this difference only reaches about 2.5%.

Next consider the conditions of large increases in permeability of the IRE cells that reflect cell death resulting from very large disruptions to the cell membrane 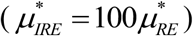; these are represented by the dashed green lines and the blue dotted lines of **Figure 5**a,b. In this scenario, the effect of IRE cell resealing plays a negligible role on the drug delivery to the RE cells for either porosity and for any field strength. The difference in drug uptake with and without IRE cell resealing never exceeded 0.6% difference in any case at any field strength).

Clearly the most influential parameters affecting the drug uptake by RE cells are (i) the magnitude of the IRE cell permeability increases, and (ii) the porosity. Comparisons for equal resealing rate constants (*τ _IRE_* = *τ _RE_*) are made between the low IRE cell permeability increase (solid black line) and the very high permeability increase (green dashed line). The difference in RE cell concentrations of these cases increases with increasing field strength. In the low porosity case the percent difference exceeded 10% (33.8 μgmL^−1^ and 30.2 μgmL^−1^) beginning at a field strength of 86 kVm^−1^; at this voltage the fraction of cells irreversibly electroporated is about 0.1. The influence of IRE permeability drastically increases at higher field strengths (and higher *f _IRE_*). In the high porosity case, the percent difference exceeded 10% (76.0 μgmL^−1^ and 68.2 μgmL^−1^) beginning at a field strength of 112 kVm^−1^; at this voltage the fraction of cells irreversibly electroporated is much higher at 0.55. At high porosity, the difference in RE cell intrinsic concentrations never exceeded 18% at any voltage while at low porosity the difference exceeds 60% beginning at voltages above 111 kVm^−1^.

To explain how the IRE cells can be so influential on the drug uptake to the RE cells, consider the uptake of the drug by the IRE cells at different field strengths (**Figure 6**a,d). The case in which the IRE cells that experience very high permeability increases 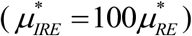 reflects the complete destruction of the cell membrane; for this scenario at low field strengths, the concentration within the IRE cells approaches the initial extracellular concentration (1000 μg mL^−1^). As the field strength is further increased, the concentrations within the IRE cells decrease asymptotically approaching the same equilibrium value as the extracellular concentrations (**Figure 6**b,e). That is because at higher field strengths the IRE cells become more permeable to the drug (recall **Figure 4**). At higher field strengths, a greater volume of tissue is converted to these high permeability IRE cells (the fraction of cells that are IRE increase with field strength as in **Figure 3**b). These two phenomena work together to lower the drug concentration in the extracellular space (**Figure 6**b,e): for high IRE cell permeability increases there is simply less drug left available in the extracellular space to be absorbed by the viable cells. The dilution of the available drug concentration in the extracellular space is more pronounced in the low porosity case (reflecting cells in tissue) than in the high porosity case (reflecting cells in dilute suspensions). Recalling the definition of porosity of Eq. (1), the definition of intrinsic drug concentration of Eq. (6) and the initial conditions (22), it is clear that the initial available drug mass is also proportional to the porosity (*m*_0_ ∝ *ɛ* C_0_) so that at lower porosity, the extracellular space is depleted more rapidly.

**Figure 6.**
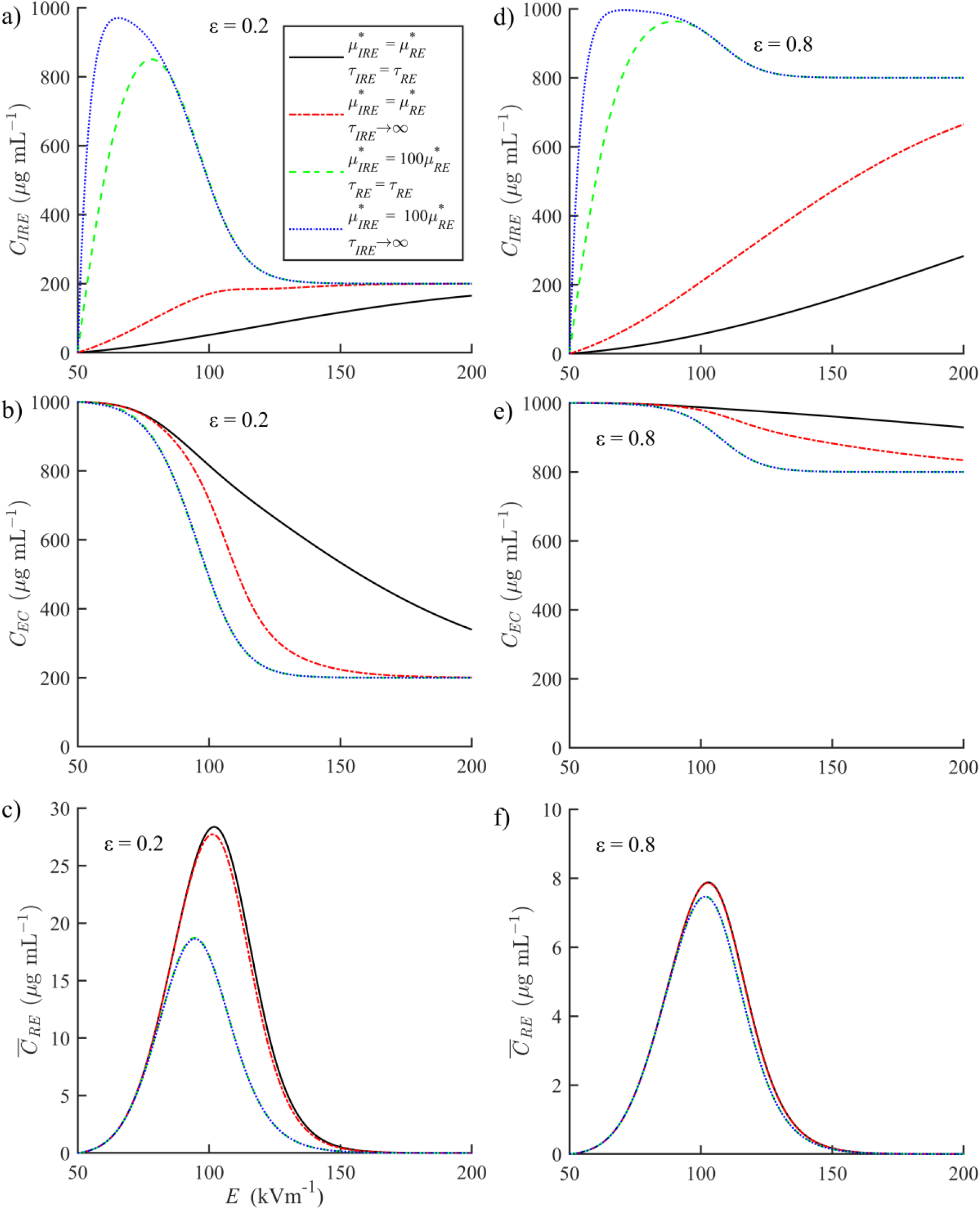
Field dependence of concentration values 400 s after exposure to 4 pulses for low porosity cases: a) intrinsic concentration in the IRE cells, b) intrinsic extracellular concentration, c) total volume averaged concentration of RE cells and high porosity cases low porosity cases: d) intrinsic concentration in the IRE cells, e) intrinsic extracellular concentration, f) total volume averaged concentration of RE cells

The intrinsic concentrations presented in **Figure 5** could be misinterpreted to imply that the total drug delivered to surviving RE cells increases with increasing field strength. As described by Eq. (8), the total drug delivered to the surviving RE cells is proportional to the total volume averaged drug concentration (which has been plotted in **Figure 6**c,f). With increasing field strength (for all cases) the drug mass delivered to RE cells increases initially, reaches a maximum, and then asymptotically approaches zero. At lower field strengths, the increases in drug mass reflect that with increasing field strength come increases in the fraction of the surviving cells that are permeabilized (**Figure 1** and **Figure 3**a) and higher permeability increases (**Figure 4**). At higher field strengths the decreases in drug mass reflect that with higher field strengths come decreases in the fraction of cells that survive (**Figure 1** and **Figure 3**b).

The influence of cell resealing on drug mass uptake is nearly negligible in all cases. In the low porosity case, the 2 order of magnitude difference in IRE cell permeability increases greatly influences the total drug delivered to the RE cells: there is a maximum difference in the total volume averaged concentration of 14.6 μgmL^−1^ that occurred at a field strength of 108 kVm^−1^. In the high porosity case its influence could be considered negligible: the 2 order of magnitude IRE cell permeability difference resulted in a maximum difference in the total volume averaged concentrations of only 0.7 μgmL^−1^ and that occurred at 113 kVm^−1^. At intermediate field strengths, the maximum total drug delivered to the surviving RE cells in the low porosity case (28.4 μgmL^−1^ at 102 kVm^−1^) case is more than 3 times higher than in the high porosity case (7.9 μgmL^−1^ at 103 kVm^−1^); this is because the volume occupied by cells in the low porosity case is 4 times higher. The concentrations in the cells uninfluenced by the electroporation have not been shown because the uptake to these cells is negligible; in all cases studied for all field strengths, the intrinsic concentrations of the NE cells never exceeded 0.05 μgmL^−1^.

When the electroporation is conducted to deliver drugs to cells, the electric field strength is chosen to be high enough to allow for significant permeability increases in the RE cells, but not so high as to result in excessive irreversible electroporation. At a field strength of 97 kVm-1 the evaluation of Eqs. (13) and (14) result in a large fraction of surviving cells that experience permeability increases (*f_RE_* = 0.84) and a low fraction of the total cells that are irreversibly electroporated (*f_IRE_* = 0.15). With this in mind Eqs. (13)–(20) are evaluated at an electric field strength of 97 kVm^−1^ for a low porosity case (Figure 7a,b,c) and for a high porosity case (Figure 7d,e,f).

**Figure 7.**
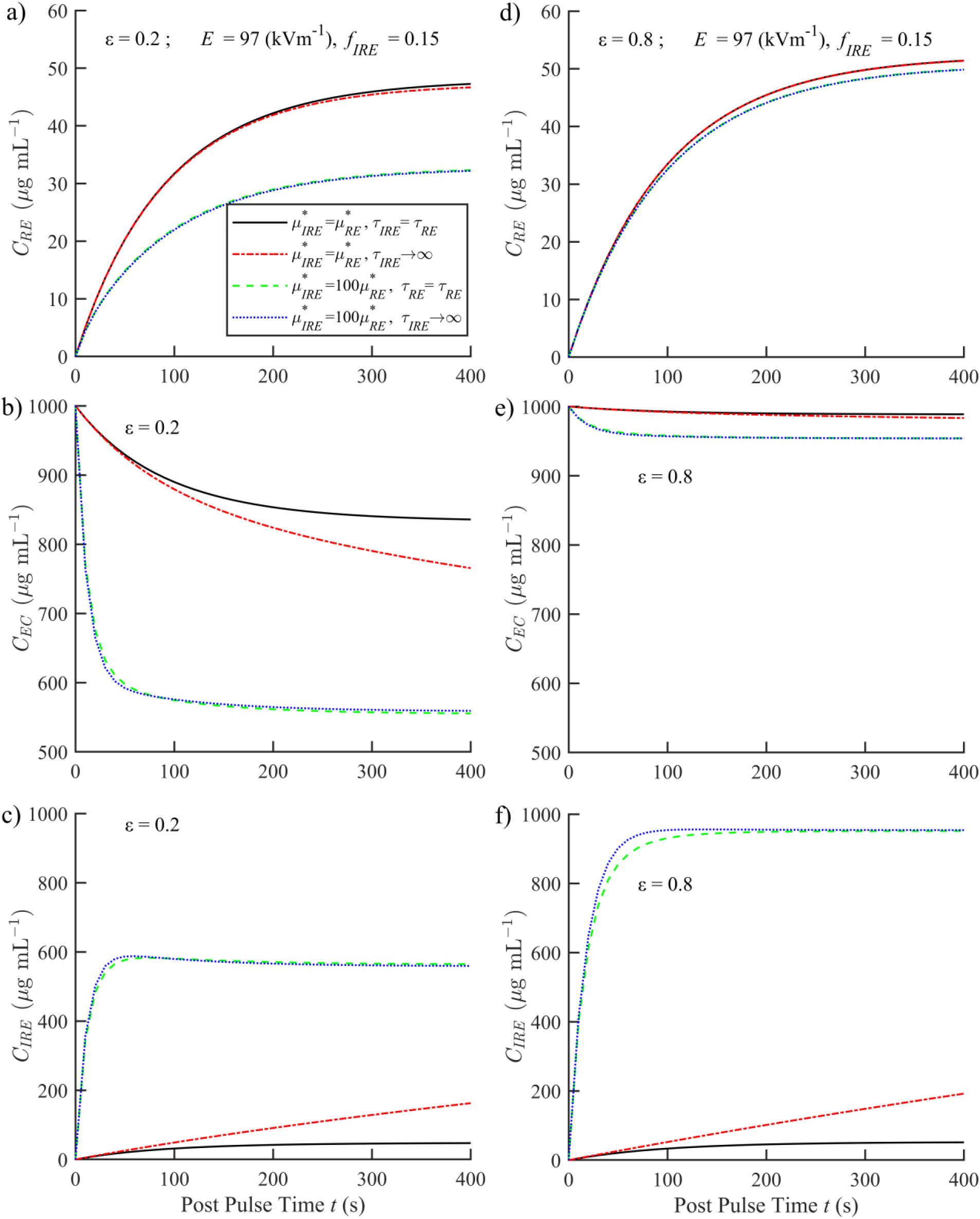
Transient intrinsic concentration values following 4 pulses at 97 kVm^−1^ which corresponds to 15% of total cells irreversibly electroporated. Concentration values of the low porosity in the space occupied by the a) reversibly electroporated cells, b) extracellular medium, c) irreversibly electroporated cells. And concentration values of the high porosity in the space occupied by the d) reversibly electroporated cells, e) extracellular medium, f) irreversibly electroporated cells.

The transient levelling off in the rates of RE cell concentration values (for all cases **Figure 7**a,d) reflect the resealing of the RE cells that is dictated by the resealing time constant *τ* _RE_. At this field strength the influence of IRE cell resealing on the uptake of drugs to the RE cells is negligible for all cases. Of these cases, the largest difference in RE cell uptake (with and without IRE cell resealing) occurs at 400 s in the low porosity case when permeability increases of IRE and RE cells are equal; here there is only about a 1% difference.

Because the rate IRE cell resealing was shown to be insignificant in RE cell drug uptake, the following analysis is made for cases with IRE resealing (*τ _IRE_* = *τ* _RE_) in comparisons between the low IRE permeability increase (the solid black line) and the high permeability increase (green dashed line). The effect of IRE cell permeability on RE cell drug uptake is much less pronounced in the high porosity case (Figure 7d); even with the 2 order of magnitude difference in IRE cell permeability; at 400 s the RE cell intrinsic drug concentrations differ by less than 3% (50.6 μg mL^−1^ vs 49.2 μg mL^−1^). When the porosity is low the variation in drug uptake by RE cells is much more pronounced (Figure 7a); at 400 s after the final pulse the RE concentrations in the high IRE permeability increase is 32.4 μg mL^−1^ while that of the low permeability increase is much higher at 46.6 μg mL^−1^.

The explanation of these transient results at moderate field strengths is as follows. The rate of drug uptake and the magnitude of drug uptake (to any cell type) are proportional to the concentration difference between the intracellular space and the extracellular space and the magnitude of the mass transfer coefficient (Eq. (21)). When the IRE cell permeability increase is high, the extracellular concentrations (**Figure 7**b,e) rapidly approach equilibrium with the RE cell concentrations (**Figure 7**c,f). Because the total available drug mass is proportional to the porosity, this equilibrium value in the low porosity case is lower than that of the high porosity case. Again the concept is that when the IRE cell permeability is very high and the porosity is low, the uptake of the drug by the IRE cells rapidly depletes the EC of the drug; this leaves less drug available for the RE cells and thus there is a lower RE cell drug concentration compared to the cases of either lower IRE permeability increases or in conditions of higher porosity.

## 4 CONCLUSIONS

This study considers the modelling of post pulse drug uptake by cells in electroporated tissue. A theoretical compartmental representation is used to distinguish the conservation of drug in each of the volumes: the extracellular space, the space occupied by reversibly electroporated cells, the space occupied by irreversibly electroporated cells, and the space occupied by cells unaffected by the electric field. The fraction of the total volume of each of these domains is related to the magnitude of the electric field using functions that are curve fit from published experimental data. To estimate the permeability increases of reversibly electroporated cells, the fraction of the cell wall occupied by long lasting electropores is related to the magnitude of the local electric field in a parabolic function that is determined from published experimental data.

A parametric investigation is conducted to determine the conditions under which the presence of irreversibly electroporated cells influences the post pulse uptake of drug by reversibly electroporated cells. The permeability increases and the resealing rate constant of irreversibly electroporated cells were chosen to reflect two types of IRE cell death. Cell death resulting from the complete and permanent destruction of the cell membrane is represented by a permeability increase two orders of magnitude higher than that of reversibly electroporated cells and a very large magnitude resealing time constant. Cell death resulting from secondary effects associated with moderate and reversible electropores is represented by a permeability increases and a resealing time constant which are equal to those of reversibly electroporated cell.

The resealing of irreversibly electroporated cells is shown to have minor influence on RE cell drug uptake and this is noticeable only when the permeability increase of the irreversibly electroporated cells is high in the low porosity case at high field strengths (where the fraction of all cells irreversibly electroporated is high). The drug uptake by reversibly electroporated cells is much more sensitive to the magnitude of the permeability increase of the irreversibly electroporated cells. This influence increases with electric field strength and decreases with porosity. Below electric field magnitudes associated with moderate cell death (‘15%) only in the low porosity case is this influence significant. \

The physical explanation behind these findings are that the uptake of the drug by the reversibly electroporated cells is reduced when the extracellular drug concentration is decreased. The drug mass in the extracellular space is rapidly depleted when both the volume occupied by irreversibly electroporated cells is a high (relative to the extracellular volume) and the permeability of these irreversibly electroporated cells is high. Both these conditions occur at higher electric field strengths and lower porosity.

The findings of this study show that the uncertainty that exists in the magnitude of the permeability increases of irreversibly electroporated cells is probably unimportant in applications of cells in high porosity dilute suspensions. In applications of electroporation of cells in low porosity tissues, the uncertainty of irreversibly electroporated cells should be considered in regions of tissue experiencing field strengths for which the fraction of cells irreversibly electroporated exceeds 0.1.

## 5 Appendices

## 5.1 Appendix I:Derivation of the Four Equation Model

Consider an elementary control volume (CV) of dimensions Δ*x* ×Δ*y* ×Δ*z* (here the total volume is V*_T_* = Δ*x*Δ*y*Δ*z*) as depicted in **Figure 8**. Under the approximation of uniform porosity, the opposite faces of the control volume have equal areas occupied by the extracellular space. On this control volume, the area in the *y-z* plane that is occupied by the *EC*, *A_EC_*, does not change with position, *x*, or in time, *t*.

**Figure 8:**
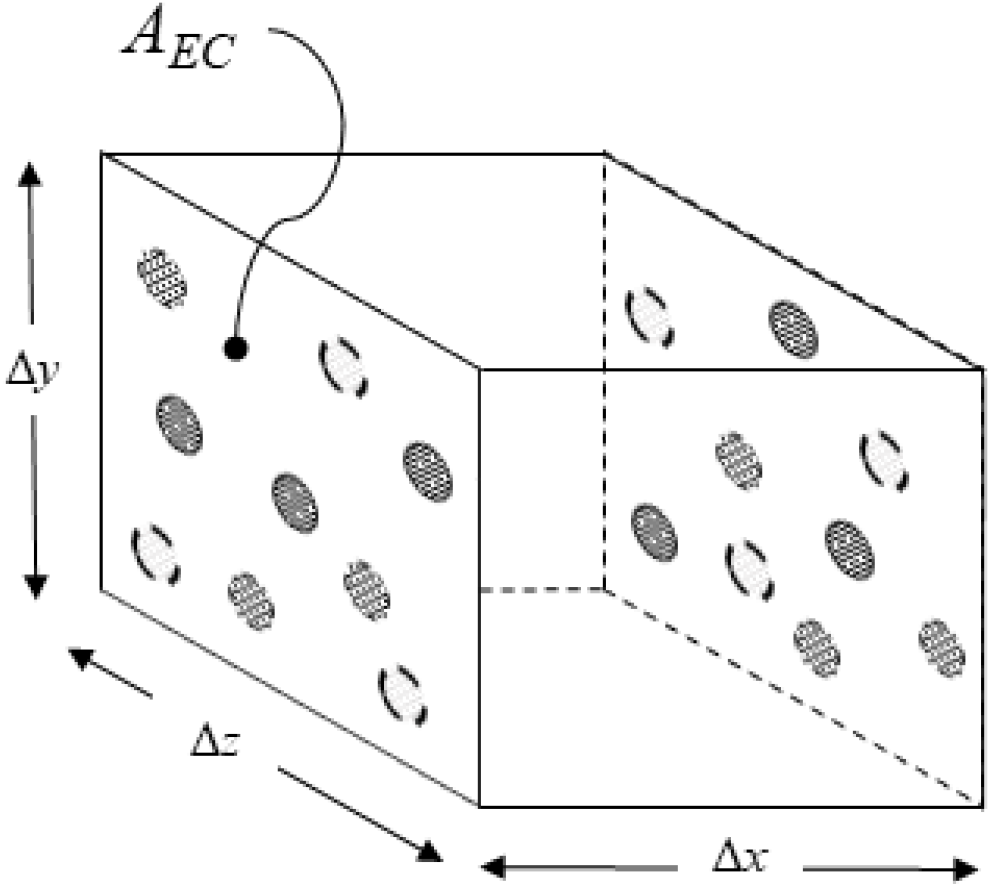
A representative control volume of dimensions Δ*x*×Δ*y*×Δ*z* with 3 different cell types.

All mass transfer to/from any cell type occurs through an exchange between that cell and the extracellular matrix. Following the electroporation pulse, and at the microscopic level, this is modelled in as a Fickian process *across the cell membrane* so that transmembrane transport is proportional to the difference in the intrinsic drug concentrations on either side of the cell wall. The membranes of the different cell types may have different resistances to mass transfer. Additionally within this CV, the rate of mass uptake by each cell type is also proportional to the total number of cells of that type (volume occupied by cells of that type within the CV). In this representation, the rate of total mass uptake by each cell type is proportional to (i) the concentration difference across the cell membrane, (ii) the resistance of the cell membrane to mass transfer, and (iii) the total number of cells (volume of cells) in the CV. With that in mind and following the derivations presented in [8] [18] [17], the transport of mass into each of the cell domains are represented as follows. The rate of mass uptake by non surviving IRE cells:

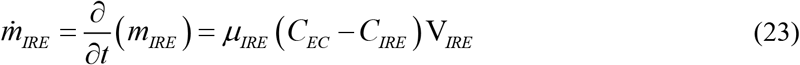

The rate of mass uptake by surviving RE cells:

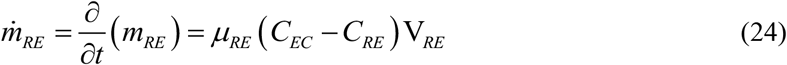

The rate of mass uptake by surviving NE Cells:

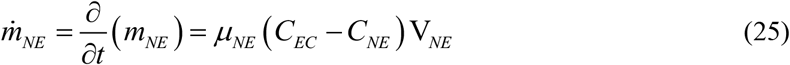

The rate change in mass storage within the extracellular space (*m_EC_*) is decreased by the net outward flux to the neighboring CV’s and is reduced with mass uptake into each of the three cell types (IRE, RE, and NE).

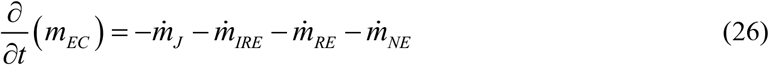

Here 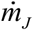 is the net mass transfer across the CV boundaries and could be a result of diffusion and of advection by interstitial flow velocity v. Here the 1-d problem is considered, though the derivation may be extended to multi-dimensions. In this case (and referring to Fig. (2)) the area of the *y-z* planes of the control volume that are made up of EC space is *A_EC_*. Within the EC, the net rate of mass flux leaving the CV is related to the rate of mass transfer per unit area, 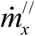 by the relation

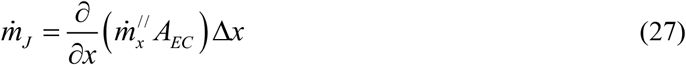

Here the rate of mass flux is governed by:

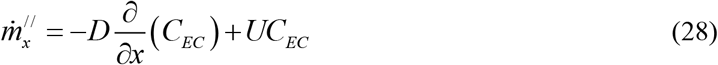

Here *D* is the effective diffusion coefficient and *U* is the average seepage velocity normal to the *y-z* planes of the CV boundaries. Substituting (28) into (27) and assuming a constant porosity and rearranging yields:

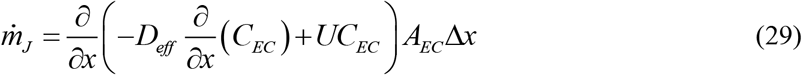

Noting that for this CV, *A_EC_* Δ*x* = V_*EC*_ and substituting in (29) and (23)–(25) into (26) results in:

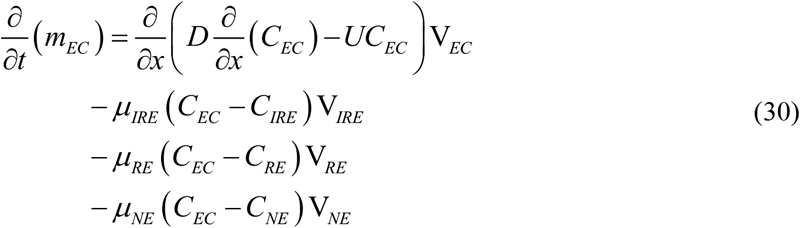

Substituting the definitions of the intrinsic concentrations of Eq. (6) and the relations of the different volumes of Eq. (5) into Eqs. (30) and (23)–(25) and dividing by total volume results in the coupled set:

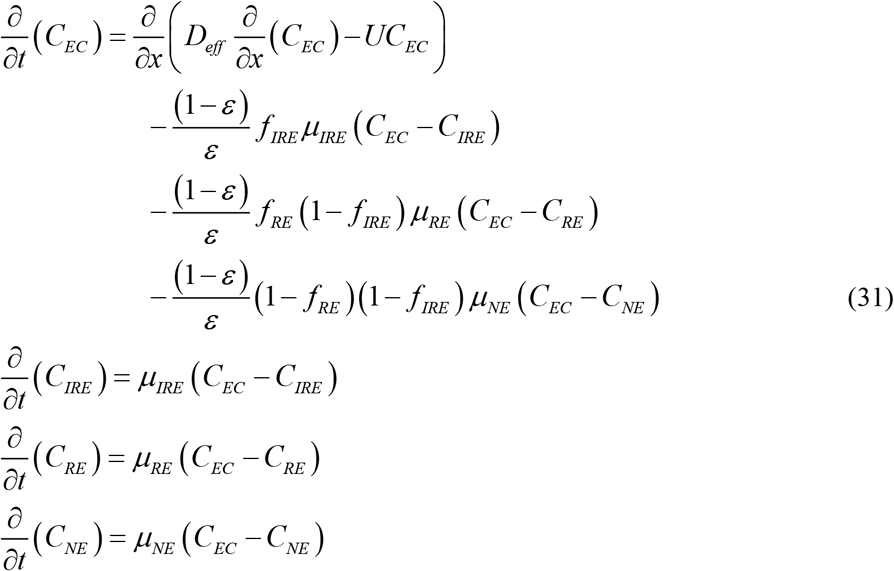

Note that the four equation representation of Eqs. (31) is immediately reduced if the mass transfer coefficient of the non-electroporated cells approaches zero. In many cases this would be a reasonable approximation (it is after all the very low permeability to drug uptake of the unaltered cell wall that usually motivates the application of electroporation). In most practical applications the electric field is spatially inhomogeneous so that the field dependent transport coefficients, *μ_RE_* and *μ_IRE_*, and the field dependent volume fractions, *f_IRE_*, *f _RE_*, and *f _NE_*, will vary by position..

This study conducts parametric investigations to determine under what conditions the IRE cells influence the drug delivery to the RE cells. The study of this paper considers the case in which there are no spatial variation in any parameter value so that the diffusive and advective terms may be ignored and that the governing equations simplify to the set of coupled reaction ODE’s of Eqs. (9)–(12).

## 5.2 Appendix II: Verification of the Numerical Simulation

Due to the nonlinearity introduced by the exponential decay terms in the set Eqs. (21), an exact analytical solution to the Eqs. (21) was not found and the results presented were determined numerically. To verify the accuracy of the numerical methodology, a simplified set of equations is solved here and a comparison with the exact solution to this simplified system is presented. The reduced model that is presented here corresponds to the case when *μ_NE_* = 0, *τ _RE_* → ∞, and *τ _IRE_* → ∞ so that the set of ODEs is represented by the three equations:

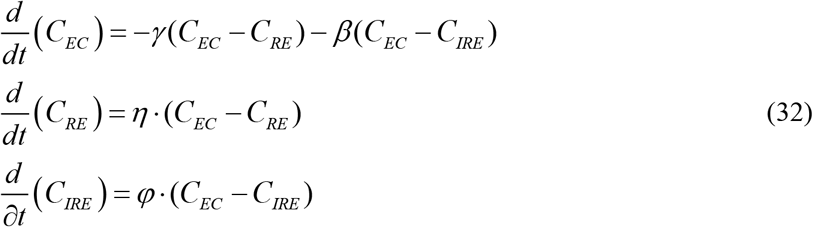

Here the coefficients of this much simplified case are related to those presented in this study by:

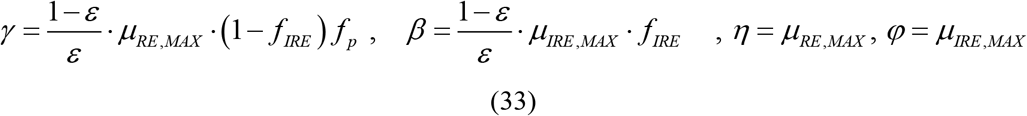

The system is solved subject to the initial conditions

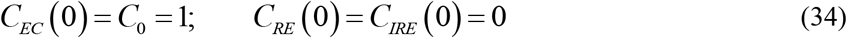

The solution to this problem may be determined using the Laplace Transform method so that:

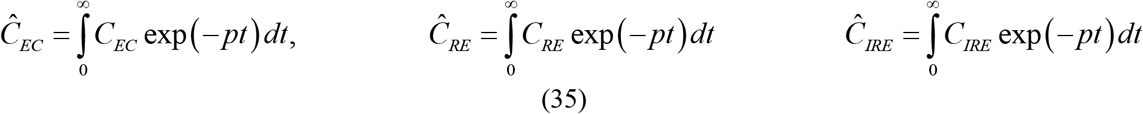

So that (32) may be represented by:

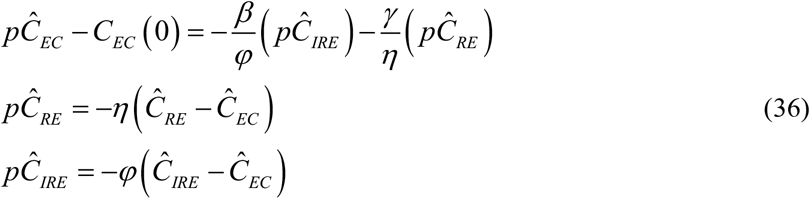

Noting that the second and third expressions of (36) can be rewritten:

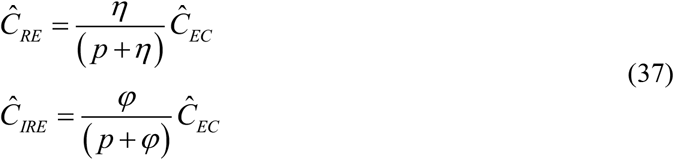

Following the substitution of (37) and algebraic manipulation, the first term of (36) may be rewritten as:

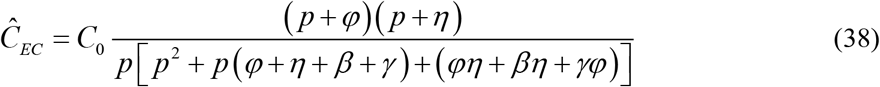

A description of the development of the inverse transform of (38) is detailed in section 2.43 (ii) on pages 21-22 of the book [33]. Essentially if a transform of *y* (*p*) has the form:

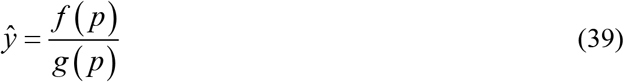

Where *f*(*p*) and *g*(*p*) are polynomials in *p* that have no common factor, the degree of the numerator, *f*(*p*), being lower than that of the denominator, *g*(*p*), and if the denominator may be expressed by:

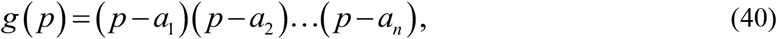

where *a*_1_, *a*_2_, … ,*a_n_* are unique constants (real or complex), then the function *y*(*t*) whose transform is given by 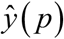 is represented by:

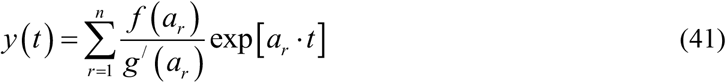

here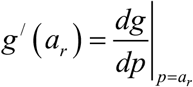.

So that applying (39)–(41) onto Eq. (38):

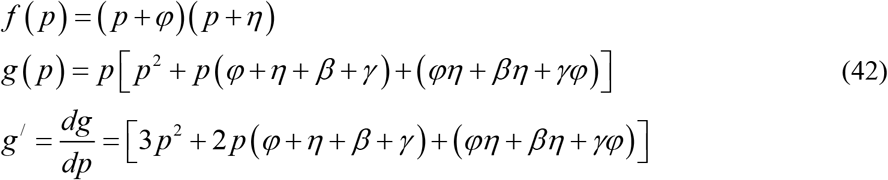

The inverse transport of (41) is evaluated at the roots of the denominator of (39) which corresponds to the roots of:

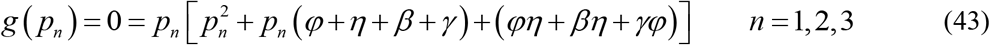

In this case the eigenvalues corresponding to the three roots of (43) are:

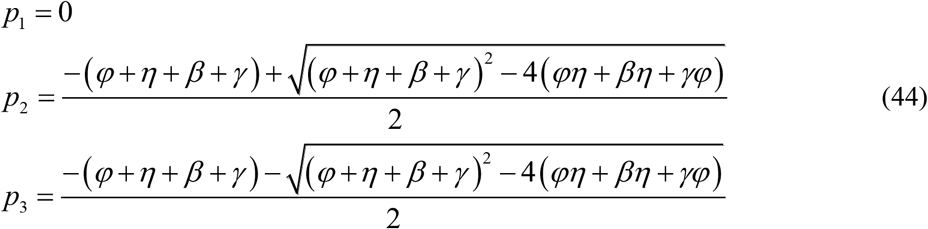

In this way, the inverse transform of (38) is represented by:

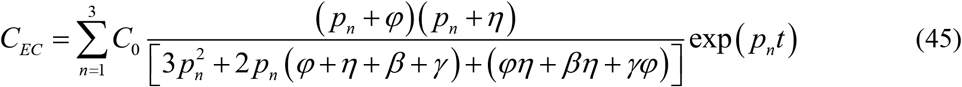

And applying the relations of Eq. (37) to Eq. (45), the concentrations within the cellular regions:

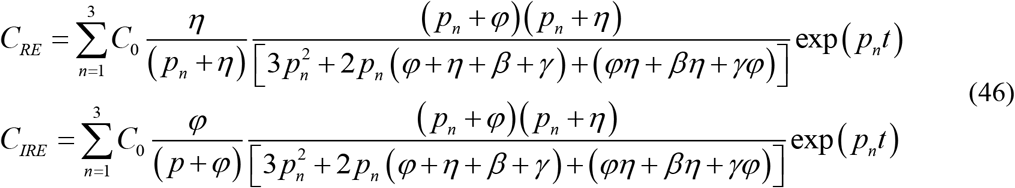

In the comparison between the numerical and exact solution the numerical solution is evaluated using the parameter values: *C*_0_ = 1 mg/mL, *ɛ* = 0.2, *μ_RE,MAX_* = 5×10^−5^ s^−1^, and*μ_IRE,MAX_* =1×10^−3^ s^−1^ along with the prediction of Eqs (13) and (14) at an electric field magnitude of 1.0 kV/cm which estimates the volume fractions, *f_IRE_* = 0.2144 and *f_P_* = 0.8769. These parameter values correspond to the exact solution parameter values *γ* = 1.378×10^−4^ s^−1^, *β* = 8.575×10^−4^ s^−1^, *η* = 5×10^−5^ s^−1^, and *φ* = 1×10^−3^ s^−1^ so that the exact solution will have the corresponding eigenvalues: *p*_1_ = 0, *p*_2_ = −1.92552×10^−3^, and *p*_3_ = −1.19786×10^−4^.

With these values a direct comparison between the numerical and analytical solution to the intrinsic concentrations in the remaining domains has been made. The instantaneous percent error in the three domains is plotted in **Figure 9** are reasonable.

**Figure 9.**
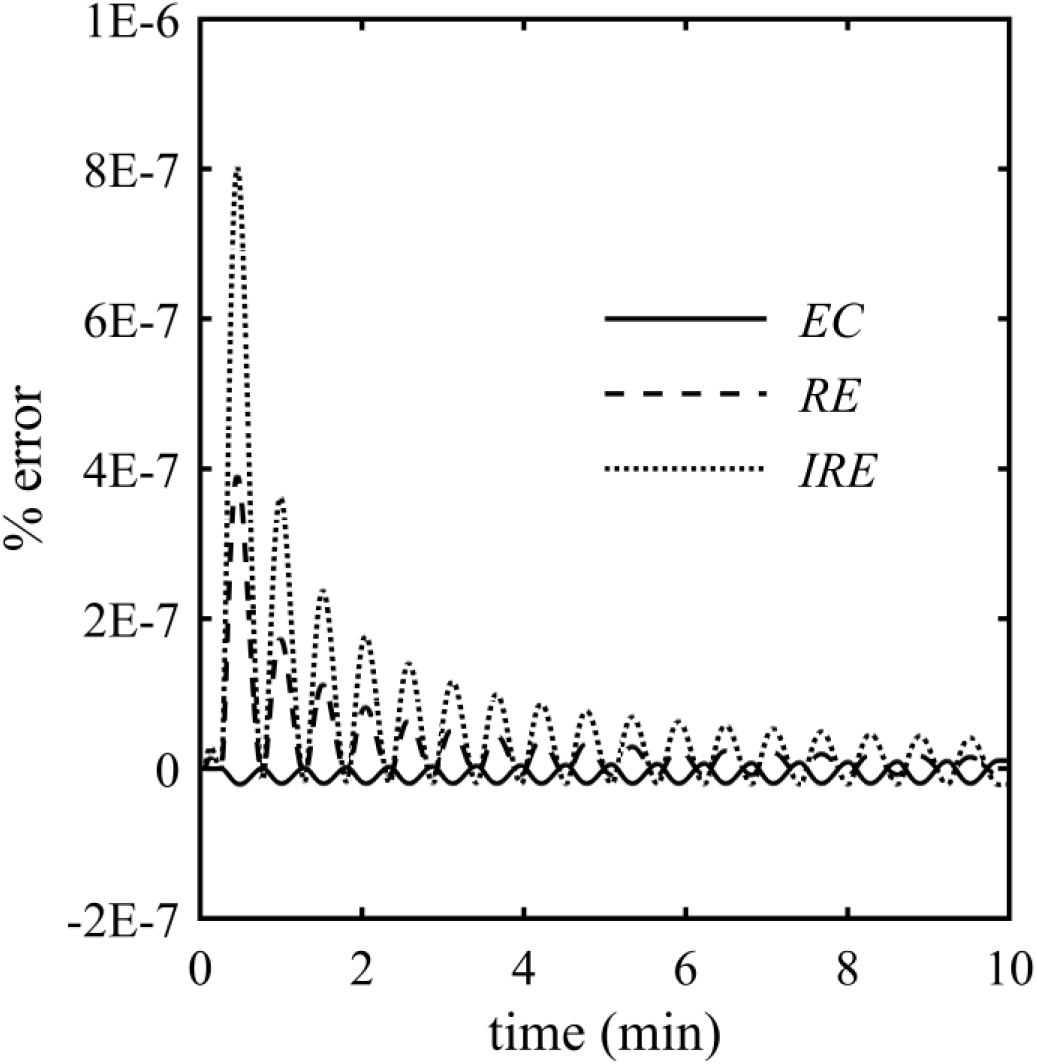
Instantaneous percent error of the numerical solution of Eq. (32) compared to the exact solution Eqs. (45)–(46)

